# Membrane remodeling by the lytic fragment of sticholysin II: implications for the toroidal pore model

**DOI:** 10.1101/620336

**Authors:** H. Mesa-Galloso, P.A. Valiente, R.F. Epand, M.E. Lanio, R.M. Epand, C. Alvarez, D.P. Tieleman, U. Ros

## Abstract

Sticholysins are pore-forming toxins of biomedical interest and represent a prototype of proteins acting through the formation of protein-lipid or toroidal pores. Peptides spanning the N-terminus of sticholysins can mimic their permeabilizing activity and together with the full-length toxins have been used as a tool to understand the mechanism of pore formation in membranes. However, the lytic mechanism of these peptides and the lipid shape modulating their activity are not completely clear. In this paper, we combine molecular dynamics (MD) simulations and experimental biophysical tools to dissect different aspects of the pore-forming mechanism of StII_1-30_, a peptide derived from the N-terminus of sticholysin II. With this combined approach, membrane curvature induction and flip-flop movement of the lipids were identified as two important membrane remodeling steps mediated by StII_1-30_-pore forming activity. Pore-formation by this peptide was enhanced by the presence of the negatively-curved lipid phosphatidylethanolamine (PE) in membranes. This lipid emerged not only as a facilitator of membrane interactions but also as a structural element of the StII_1-30_-pore that is recruited to the pore ring upon its assembly. Collectively, these new findings support a toroidal model for the architecture of the pore formed by this peptide and provide new molecular insight into the role of PE as a membrane component that easily accommodates into the ring of toroidal pores aiding in its stabilization. This study contributes to a better understanding of the molecular mechanism underlying the permeabilizing activity of StII_1-30_ and peptides or proteins acting via a toroidal pore mechanism and offers an informative framework for the optimization of the biomedical application of this and similar molecules.

**State of significance:** We provide evidence about the ability of StII_1-30_ to form toroidal pores. Due to pore assembly, StII_1-30_-pore induces membrane curvature and facilitates flip-flop movement of the lipids. The negatively-curved lipid PE relocates from the membrane into the pore ring, being also a structural element of the pore StII_1-30_ forms. This peptide emerged as a new tool, together with the full-length toxin, to understand the mechanism of toroidal pore formation in membranes. This study provides new molecular insight into the role of curved lipids as co-factors of toroidal pores, which could aid in its stabilization by easily accommodating into the ring. This framework could underpin strategies for the rational use of peptides or proteins acting via toroidal pores.

## I. Introduction

Permeabilization of lipid membranes by pore-forming toxins (PFTs) has received special attention due to their role as potential virulence factors. These proteins bind and oligomerize in membranes, leading to membrane permeabilization and cell death (1–3). Therefore, they are also considered useful tools to study the basic molecular mechanisms of protein insertion into membranes. These toxins show a dynamic interplay with lipid membranes since they require a large conformational change for membrane insertion. In turn, they strongly modify the membrane structure upon interaction with the bilayer (4, 5). PFTs are usually classified according to the structure of the membrane-integrated domain as α- or β-PFTs (6). These lytic fragments commonly show structural homology with natural membranolytic peptides such as antimicrobial peptides (AMPs), which are typically molecules in which the pore-forming domain covers the entire length of the polypeptide (7). Based on this homology, synthetic peptides have been used as a valuable tool to study the contribution of lytic fragments to the activity of some pore-forming proteins such as the pro-apoptotic protein Bax (8) and actinoporins (9–13).

Actinoporins are a unique class of highly hemolytic eukaryotic α-PFTs, exclusively found in sea anemones, whose putative receptor is sphingomyelin (SM) (14–17). Lipid mixing from different phases (18) and induction of membrane curvature and lipid flip-flop (15, 19) are some membrane remodelling effects of these toxins, which are closely related to their pore-forming function. Indeed, these effects on membrane structure are commonly associated with their ability to form hybrid protein-lipid or toroidal pores (15, 19). It has been widely accepted that the N-terminal α-helix of actinoporins is the protein fragment that builds the pore, which is also lined by phospholipids head groups (15, 19, 20). In this regard, actinoporins represent a prototype of proteins acting through the formation of protein-lipid pores. Together with other virulence factors such as the cholesterol dependent cytolysins (CDCs) and the apoptotic proteins Bax and Bak, they have been widely used as a tool to understand the mechanism of toroidal pore formation in membranes (4, 5, 21, 22).

Sticholysins I and II (StI and II) are actinoporins produced by the Caribbean Sea anemone *Stichodactyla helianthus* (23). These toxins are of current interest as an active component of a vaccine platform (24, 25) and of immunotoxin constructs active against tumor cells (26, 27). Peptides reproducing the N-terminus of StI sequence 1-31 (StI_1-31_) and StII sequence 1-30 (StII_1-30_) (Figure 1A) can mimic the permeabilizing ability of these toxins in red blood cells and liposomes (10, 12, 13). In particular, StII_1-30_ is the most active peptide (12, 13, 28). Its forms pores in human red blood cells with a similar size to those formed by the full-length toxin (9). StII_1-30_ has emerged as a good candidate to replace the whole toxins in some biomedical applications such as an active component of a vaccinal platform to promote the delivery of different molecules to the cytosol and enhance the immune response in anti-cancer therapies (24, 25). Previously, we have analysed the conformational properties of StI_1-31_ and StII_1-30_ in solution and in membrane mimetic systems and their ability to permeabilize lipid vesicles (9, 12, 13, 28, 29). In solution, StII_1-30_ acquires an α-helical fold in coiled-coil self-associated structures, without requiring membrane assistance (29). Based on these findings, we previously suggested a model in which StII_1-30_ membrane permeabilizing activity is favoured by peptide pre-association in solution. However, the lytic mechanism of StII_1-30_ is not completely clear. In particular, the interplay between lipid shape and StII_1-30_-induced lipid rearrangements has not been explored yet (30). Moreover, there is no current information about the structure of the pores that this peptide forms in lipid membranes.

**Figure 1.**
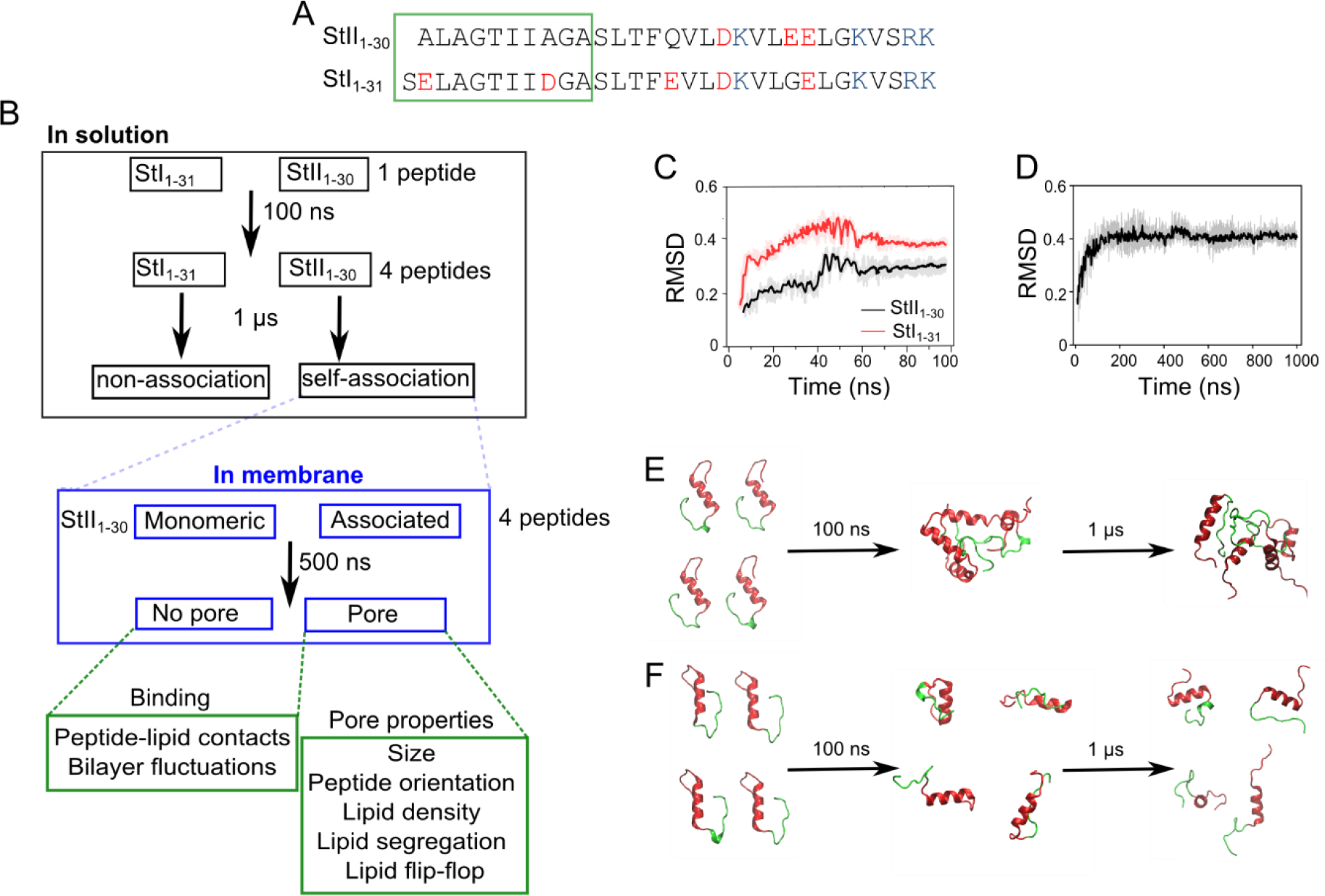
StII_1-30_ has a high propensity to associate in solution. **(A)** Sequences of StII_1-30_ and StI_1-31_. Acidic amino acid residues are in red and basic amino acid residues are in blue. The 1-10 (StII_1-30_) and 1-11 (StI_1-31_) regions are highlighted in green. **(B)** Workflow of the simulations and analysis. Snapshots of the simulation box of StII_1-30_ **(C)** RMSD calculated during equilibration of monomeric peptides StII_1-30_ and StII_1-31_ in solution. **(D)** RMSD calculated for the association of StII_1-30_ in solution. Snapshots of the MD obtained with StII_1-30_ **(E)** and StI_1-31_ **(F)** at time 0, 100 and 1 μs of simulation in solution. Each peptide is represented in two colors: the hydrophobic sequence 1-10 (StII_1-30_) or 1-11 (StI_1-31_) regions are in green and the 11-30 (StII_1-30_) or 12-31 (StI_1-31_) regions are in red.

In this study, we characterized in atomistic detail the mechanism of membrane perturbation leading to pore formation by StII_1-30_. Using molecular dynamics (MD) simulations, we modelled membrane reshaping and pore formation in membranes including phosphatidylcholine (PC), SM and phosphatidylethanolamine (PE). From our simulations, we predicted several structural properties of the pore including size, peptide topology and lipid segregation into the pore, the ability of the peptide to induce membrane curvature and flip-flop movement of the lipids, and the key role of PE in pore-formation. In vitro, we focused on the effect of PE on StII_1-30_ binding to lipid monolayers and liposomes and on its permeabilizing activity in vesicles. Moreover, we also assessed the capability of StII_1-30_ to induce membrane curvature and lipid flip-flop in these membrane mimetic systems. Our data identified PE as a critical membrane component that favours StII_1-30_ activity by modulating membrane properties required for more efficient binding and pore formation and also as a lipid constituent of the pore. Membrane curvature and lipid flip-flop induction emerged as key steps in the mechanism of membrane permeabilization by this peptide. This work provides the first evidence supporting toroidal pore formation in the lytic action of StII_1-30_ and highlights a structural role of lipid shape on the pore formed by this peptide.

## II. Materials and Methods

### II.1 Chemicals and reagents

Lipids:1-palmitoyl-2-oleoyl-glycero-3-phosphocholine (POPC), porcine sphingomyelin (SM), 1-palmitoyl-2-oleoyl-glycero-3-phosphoethanolamine (POPE), 1,2-dipalmitoleoyl-phosphatidylethanolamine (DiPoPE) and pyrene-labeled PC (pyPC) were purchased from Avanti Polar Lipids (Alabaster, AL, USA) with 99% purity and used without further purification. The peptides were produced by solid-phase synthesis and further purified to >96% purity by reversed-phase HPLC by GL Biochem (Shanghai, China). Peptide concentration was evaluated by a Micro BCA protein assay kit provided by Pierce (Illinois, USA).

### II.2 MD simulations

An overview of our simulations is shown in figure 1 (Figure 1B). Peptide systems were created in an 8 × 8 × 8 nm water box in solution or 12 × 12 × 12 nm in membranes. First, the structures of StII_1-30_ and StI_1-31_ were obtained from the structure of StII (PDB 1O72) (31) and StI (PDB 2KS4) (32) respectively, and equilibrated in solution for 100 ns. Then, four peptides in solution were simulated during 1 μs to allow peptide self-association. All systems in solution were created with a distance of 2 nm in the *x*-dimension and 4 nm in the *y*- and *z*- dimensions between each peptide in an 8 × 8 × 8 nm water box. Membrane systems were created with four peptides of StII_1-30_ starting from two different initial configurations: i) monomeric or non-self-associated or ii) pre-associated in solution. Each peptide system was simulated in the presence of lipid membranes of different compositions: POPC:SM (50:50) and POPC:SM:POPE (50:40:10) (Table 1) using a box dimension of 12 × 12 × 12 nm. Lipid systems contained 240 lipid molecules, which corresponded to a peptide/lipid (P/L) ratio of approximately 1/60. Peptide molecules were added to one side of the membrane, mimicking in vitro experiments in which peptide molecules are initially added to the external monolayer of liposomes. Membrane-peptide systems were generated using CHARMM-GUI and additional in-house developed software (33). The system was solvated using the TIP3 water model, and sodium and chloride ions were added to neutralize the system and to reach 150 mM final concentration. Each system was initially energy-minimized for 50 000 steps. After minimizing its energy, the system was equilibrated for 150 ps before a 4-ns MD simulation at constant pressure (P=1 atm, semi-isotropic coupling) and temperature (T= 310 K). Periodic boundary conditions were applied in all three dimensions. Berendsen coupling protocols were used for the first 150 ps of the equilibration simulations (34, 35) and Nose-Hoover (36) and Parrinello-Rahman protocols were used for temperature and pressure coupling in the 4ns of equilibration and the production simulation (37). Long-range electrostatic interactions were computed using the Particle Mesh Ewald (PME) method (38, 39). Lennard-Jones energies were truncated at 1.2 nm. Bond lengths were constrained with LINCS (40). Following this initial setup, each system was simulated for 500 ns, at 310 K, with a time step of 2 fs. Three simulations were performed for each system, starting from different initial Maxwell distributions of random velocities. Frames were saved every 400 ps. Simulations were carried out using GROMACS version 5.0.2 (41). The force field employed was CHARMM 36 (42).

**Table 1:** Overview of the StII_1-30_-membrane simulations.

| Composition | No. of peptides | Associated peptides | Simulation Length (ns) | Replicates/Number of simulations |
| --- | --- | --- | --- | --- |
| POPC:SM (50:50) | 4 | No | 500 | 3 |
| POPC:SM (50:50) | 4 | Yes | 500 | 3 |
| POPC:SM:POPE (50:40:10) | 4 | No | 500 | 3 |
| POPC:SM:POPE (50:40:10) | 4 | Yes | 500 | 3 |

### II.3 Analysis of the MD simulations

#### Lipid-peptide interactions and bilayer alterations

Peptide-lipid contacts were defined when the phosphodiester group of a given lipid (PC, SM, or PE) was found within 5 Å from the center of mass of each individual peptide. For a given frame, the total number of contacts was calculated as the sum of all the contacts established between phosphate groups of an individual lipid molecule and the amino acid residues of the peptides following the procedure described by (43). The calculation was performed using the g_select tool implemented in GROMACS. Membrane thickness fluctuation in the process of pore formation was calculated in the region defined by the pore center using a cut-off of 1.5 nm, as described in (44). As the shape of the pore is non-uniform across the membrane, the central region of the pore was defined as ± 0.7 nm on each side of the center of the bilayer. The thickness of the membrane around the pore region was calculated using frames every 100 ns as the average phosphate-phosphate distance of opposite monolayers, at each time point. Fluctuations in the thickness were calculated with respect to the initial simulation time.

#### Pore size

The presence of a transmembrane pore was determined following a clustering method as described in (44). Here only the atomic coordinates of the phosphate atoms of POPC, POPE, and SM were analyzed. For each lipid, the phosphate atoms of each monolayer were considered as two independent clusters and pores were identified when the phosphate groups of the lipids formed only one cluster. For the cluster analysis, a minimal distance of 1.0 nm between PO_4_ groups that form the upper and lower monolayer was established as a cut-off for pore definition. To estimate the size of the pore, the diameter of the circle fitting the lateral projected density formed at the lipid-water interface of the pore was calculated as described in (44) using frames every 10 ns.

#### Peptide inclination and orientation

The variation of the tilt angle was calculated over the simulation time for each peptide using the *z*-axis perpendicular to the bilayer as a reference. Peptide orientation was estimated based on the distance of the *z*-coordinate of amino acid residues located in three different regions of each peptide molecule (N-terminus: Ala1; center of mass: Gln15; and C-terminus: Lys30). The distance between the peptides and the center of the bilayer (*z*’=0) were averaged during the last 100 ns of simulation.

#### Lipid contribution to the pore

Lipid density profiles were estimated as described in (45). In brief, the plane of the bilayer (*xy*) was divided into 10 000 grid cells of ca. 0.2 nm × 0.2 nm and the number of PO_4_ groups of each lipid type in each grid was calculated. Lipid densities were averaged over the last 100 ns of simulation corresponding to the time point in which the number of each lipid type surrounding the peptides within 5 Å was constant, and the pore reached its maximum size. For each lipid type, lipid density values were normalized by considering the average density in the whole system.

The depletion-enrichment (DE) index of the lipids in the local environment of the pore was calculated considering a cut-off distance of 1.5 nm in the xy plane from a line going through the center of the pore and averaged during the last 100 ns of simulation time, in which the pore adopted a stable open configuration. The cut-off distance was chosen considering that the final diameter of the pore (approx. 2 nm) was reached after 300 ns and 400ns of simulation time in POPC:SM:POPE (50:40:10) and POPC:SM (50:50), respectively. The enrichment value of a lipid X (PC, SM, or PE) in the pore is defined as the ratio of its locus to its bulk concentrations following the equations (46, 47):

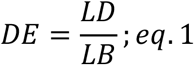

In which the lipid distribution (LD) was calculated as follows:

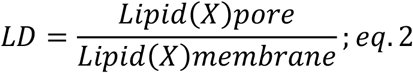

And the lipid bulk (LB) was calculated as follows:

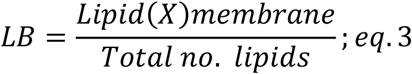

Where the total lipids are the sum of all of them in each system. The DE index was calculated for the last 100 ns of the simulation for each individual lipid type for the lower and upper monolayer together. The statistical significance was calculated by a single sample T-test, according to (47).

#### Flip-flop movement of lipids

To analyze flip-flop movement of the lipids between the two leaflets of the membrane, we consider the PO_4_ group of each lipid type, located in the pore region (1.5 nm in the x-y plane from a line going through the center of the pore). We defined the middle plane of the membrane as *z*’=0 as the mean *z*- coordinate between the lower and the upper leaflet. The upper leaflet is defined as *z*’>0 and the lower leaflet as *z*’<0. Considering this, the *z*-coordinate of the lipid phosphodiester group was calculated as a function of the simulation time.

### II.4 Binding to lipid monolayers

Surface pressure (π) measurements were carried out as previously reported (17, 48, 49) with a μThrough-S system (Kibron, Helsinki, Finland) at room temperature under constant stirring. The aqueous phase consisted of Tris-buffered saline (TBS: 145 mM NaCl, 10 mM Tris-HCl, pH 7.4). The lipid mixture (PC:SM (50:50) or PC:SM:POPE (50:45:10)) was pre-dissolved in chloroform:methanol (2:1, v:v) and was gently spread over the surface. The desired initial surface pressure (π_o_) was attained by changing the amount of lipid applied to the air-water interface. StII_1-30_ was injected into the sub-phase to achieve a concentration at which no effect on surface pressure of the air-water interface was observed. The increment in surface pressure (Δπ) was recorded as a function of the elapsed time until a stable signal was obtained.

### II.5 Preparation of lipid vesicles

Films of PC:SM (50:50) or PC:SM:POPE (50:45:10) were prepared by evaporation of stock chloroform solutions using a stream of wet nitrogen and submitted to vacuum for not less than 2 h. For permeabilizing assays, multilamellar vesicles (MLV) were obtained by subsequent hydration in the presence of 80 mM carboxyfluorescein (CF), pH 7.4 (adjusted by adding NaOH) in water and subjected to six cycles of freezing and thawing. Large unilamellar vesicles (LUV) were prepared by extruding this MLV solution through a two‐syringe LiposoFast Basic Unit extruder (Avestin Inc., Ontario, Canada) equipped with two stacked polycarbonate filters with 100 nm pore size (Nuclepore, Maidstone, UK). To remove untrapped fluorophore, vesicles were filtered through a mini‐column (Pierce, Rockford, USA) loaded with Sephadex G‐50 pre‐equilibrated with TBS. To study binding to liposomes, small unilamellar vesicles (SUV) were prepared by sonication of an MLV suspension, prepared as described above, using an ultrasonicator (Branson 450, Danbury, USA) equipped with a titanium tip and subjected to 15 cycles of 2 min sonication with intervals of 1 min rest. Titanium particles released from the probe were removed by further centrifugation at 10,000g for 10 min at 22°C. Phospholipid concentration was measured by determining inorganic phosphate according to Rouser et al (50). Lipid concentration used are given in the text.

### II.6 Binding to liposomes

Binding of StII_1-30_ to SUVs was followed by the increase in Phe or Trp fluorescence. Since StII_1-30_ does not contain any Trp in its primary sequence, the Trp-analogue peptide StII_1-30L2W_ was employed for Trp measurements (28). Fluorescence measurements were carried out at room temperature in a RF-5301PC spectrofluorimeter (Shimadzu, Tokyo, Japan) using quartz cuvettes with excitation and emission slits of 5 and 10 nm, respectively. Peptide samples were excited at λ_exc_ 240 (Phe) or 280 (Trp) nm and the emission spectra were recorded from λ 300 nm to 440 nm. The spectral correction was made by subtracting spectra measured under identical conditions but without peptide.

### II.7 Differential scanning calorimetry

The effect of StII_1-30_ and StII on the lamellar (L) to inverted hexagonal (H_II_) transition temperature of DiPoPE was assessed by differential scanning calorimetry (DSC). Lipid films were made by dissolving appropriate amounts of lipid in chloroform/methanol 2:1 (v/v) followed by solvent evaporation under a stream of nitrogen to deposit the lipid as a film on the walls of a tube. The tube was placed in a vacuum chamber for at least 2 h, to remove possible traces of solvent. Films were hydrated with 20 mM PIPES buffer (1 mM EDTA, 150 mM NaCl, 0.002% NaN_3_, pH 7.4) in the presence or not of an appropriate amount of protein or peptide and vortexed extensively to make MLVs. Calorimetric scans were carried out on a MicroCal VP-DSC differential scanning calorimeter (Massachusetts, USA). The reference and the sample solutions were degassed at room temperature prior to scanning. The scan rate was 1 °C.min^−1^, with a delay of 10 min between sequential scans in a series to allow thermal equilibration. The scans corresponding to lipid transitions were recorded in the presence and in the absence of peptide using PIPES buffer as the reference. The temperature of transition (*Tm*) was calculated as the maximum of heat capacity vs. temperature.

### II.8 Vesicle permeabilization

LUVs permeabilization was determined by measuring the fluorescence (λ_exc_ = 490 nm and λ_em_ = 520 nm) of released CF. Black plastic 96-well microplates (SPL, Life Sciences, Seoul, South Korea) were pretreated with 0.1 mg/ml Prionex (Pentapharm, Switzerland) which strongly reduces unspecific binding of protein and vesicles to plastic. Each well was filled with the TBS plus 10 μM of lipids. Finally, the peptide was added in a total volume of 200 μl at the concentration reported in the text. After mixing vesicles and peptides, the release of CF produced an increase in fluorescence, F (due to the dequenching of the dye into the external medium), which was resolved in time. Spontaneous leakage of the dye was negligible under these conditions. Maximum release was always obtained by adding 1 mM Triton X-100 (final concentration) and provided the fluorescence value f_total_. The fraction of fluorophore release (F in %) was calculated as follows:

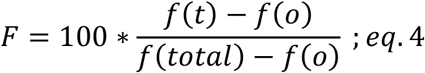

Where f(t) is the fluorescence at time t, f(total) is the maximum release upon addition of Triton X-100, and f(0) is the fluorescence before addition of the peptide. Phospholipid concentration in LUV was measured as previously reported (50).

### II.9 Measurement of flip-flop movement of the lipids

Effect of StII_1-30_ on the transbilayer movement of lipids was assessed by fluorescence spectroscopy, as described before (51). For this, LUVs were prepared as described above in TBS buffer without CF. Briefly, the asymmetrically labelled liposomes were prepared by incubating the liposome suspension (20 μM final lipid concentration) with pyrene-labeled PC (pyPC) suspension 1 μM, for 20 min at 37 °C. Changes in the fluorescence spectrum of PyPC upon peptide addition was followed in a spectrofluorimeter (Hitachi F-4500) using an λ_exc_ of 344 nm and measuring the fluorescence intensity of monomers (I_M_) and excimers (I_E_) at λ 395 and 465 nm, respectively. The time course of the transbilayer redistribution of pyPC was followed through the calculation of the I_M_/I_E_ ratio.

## III. Results

### III.1 Pre-associated in solution StII_1-30_ forms pores in lipid membranes

Previous structural studies based on low-resolution circular dichroism measurements of StII_1-30_ have shown that this peptide is able to adopt α-helical secondary structure and self-associate forming coiled-coil structures in solution without the assistance of the membrane (9, 29). Therefore, we performed atomistic MD simulations to get pre-associated StII_1-30_ in solution. The homologous sticholysin I-derived peptide (StI_1-31_) that does not self-associate was used as a control (10, 29) (Figure 1A). Knowing from previous CD studies that StII_1-30_ and StI_1-31_ have the potential to adopt similar structures to those found in the context of the protein (29), here we started the simulations from the structure that these segments adopt in their respective parental sticholysins. The peptides were further equilibrated in solution for 100 ns (Figures 1B-1E). Since monomeric peptide molecules were stable after 100 ns (Figure 1C), four peptides were further placed in a water box and simulated for 1μs to allow self-association (Figures 1D-1F). Individual StII_1-30_ molecules interacted with each other at 100 ns, forming associated structures that remained stable after 1 μs simulation (Figures 1C and 1E). Conversely, StI_1-31_ did not self-associate during 1 μs of simulation (Figure 1F).

To model pore formation by StII_1-30_ in membranes, a series of atomistic MD simulations on lipid bilayers was performed in membranes containing POPC and SM with and without POPE. SM was selected since this lipid has been proposed as the natural receptor of actinoporins (52) while POPE was selected because of its capacity to form non-lamellar structures favoring negative curvature in the membrane (53). Non-lamellar structures are essential for toroidal pore formation, where the membrane adopts both a positive curvature in the direction parallel to the pore axis, and a negative curvature in the direction perpendicular to the pore axis (54, 55).

In our simulations two configurations were modeled, each of them containing four molecules of StII_1-30_: i) non-associated StII_1-30_ and ii) pre-associated StII_1-30_. Pre-associated StII_1-30_ was obtained after 1 μs simulation (Figures 1C and 1E). Each system was simulated in three replicas. Figure 2A and 2B display representative snapshots of one of the simulations with pre-associated StII_1-30_ in DOPC:SM (50:50) or DOPC:SM:POPE (50:40:10) membranes, respectively. Statistical analysis of the replicates is summarized in Table 2. Lipid binding and membrane fluctuation took place only when StII_1-30_ started from a pre-associated conformation in solution (Figures 2C and 2D). In this condition, the increase in the average number of contacts of the peptides with the polar head of lipids in the membrane (Figure 2C) indicated membrane destabilization. Pore formation induced the increase in membrane thickness probably as a consequence of lipid recruitment to the pore region (Figure 2D). These findings are in agreement with our previous experimental data showing that pre-association in solution enhances the association of N-terminal derived peptides of sticholysins to the membrane (56).

**Figure 2.**
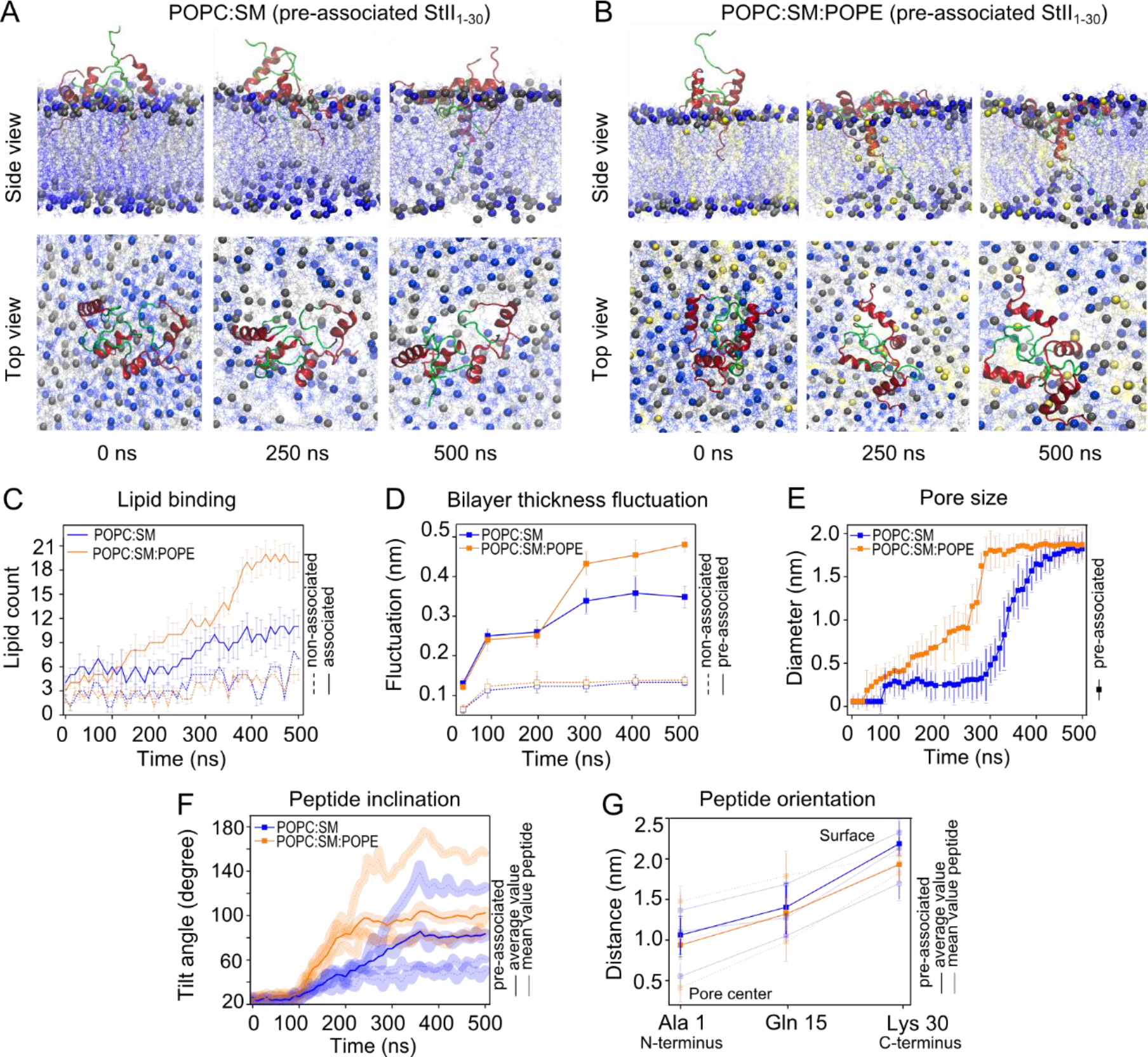
POPE favours interaction with lipids and pore formation by pre-associated StII_1-30_. Snapshots of the sequence of events in the simulations in **(A)** POPC:SM (50:50) and **(B)** POPC:SM:POPE (50:40:10). Phospholipid head groups are represented in grey (POPC), yellow (SM) or blue (PE) and peptides are represented in two colors the 1-10 segment is represented in green and the 11-30 in red. Water was removed for clarity. **(C)** Kinetics of contact average variation the four molecules of StII_1-30_ with the polar head of the phospholipids during the simulation time. **(D)** Kinetics of membrane thickness fluctuation around the pore region in the presence of StII_1-30,_ measured as the distance between two different clusters of phosphate groups located at each monolayer during the simulation time. **(E)** Increase in size kinetics of the pore formed by pre-associated StII_1-30_ in POPC membranes during the simulation time. The pore size was defined as the diameter of the circle formed at the lipid-water interface of the pore. **(F)** Kinetics of associated StII_1-30_ insertion into the bilayer, during the simulation time. The tilt angle was calculated for each peptide molecule using the *z*-axis perpendicular to the bilayer as a reference. **(G)** Orientation of associated StII_1-30_ molecules averaged over the last 100 ns of the simulation. In C-E, values correspond to the mean and the standard deviation considering the three replicates of the system. In F and G, light lines correspond to the mean values of the four peptides of each system and dark lines are the average of the three replicas.

**Table 2:** Parameters obtained from the simulations of self-associated StII_1-30_ in different lipid membrane systems.

| Parameter | Membranes |  |
| --- | --- | --- |
|  | POPC:SM | POPC:SM:PE |
| Lipid count <sup>a</sup> | 7 ± 3 | 18 ± 2* |
| Membrane thickness fluctuation (nm) <sup>b</sup> | 0.4 ± 0.03 | 0.5 ± 0.02* |
| Pore diameter (nm) <sup>c</sup> | 1.9 ± 0.4 | 2.0 ± 0.5 |
| Tilt angle (degree) <sup>d</sup> | 89 ± 12 | 98 ± 15 |
| Average distance of the N-terminus to the pore center (nm) <sup>e</sup> | 1.1 ± 0.4 | 1.0 ± 0.3 |

<sup>a</sup>Lipid count defined as lipids that contact the peptides within a distance of 5 Å from the peptides. <sup>b</sup>Membrane thickness fluctuation was calculated as the deviation of the distance phosphate-phosphate (between each monolayer) within a cutoff of 1.5 nm from the pore center. <sup>c</sup>Pore diameter was determined by the circle formed at water-lipid interface of the pore. <sup>d</sup>Tilt angle was calculated considering the angle formed between each peptide and the z axis perpendicular to membrane plane. <sup>e</sup>Distance between the center of mass of the peptide and the center of the pore, calculated as the z coordinate of the peptide respect to the center of the bilayer (z'=0) averaged during the last 100 ns of simulation. Values correspond to the mean and standard deviation from the three replicates of the system. \*indicates that the values are statically different with p values calculated from a two-tailed distribution reported with a confidence level of 0.05.

### III.2 StII_1-30_-induced pores have a toroidal shape

In our MD simulations, StII_1-30_ induced membrane curvature in both PC:SM and POPC:SM:POPE systems, as a consequence of pore formation (Figures 2A and 2B). Moreover, our simulations showed that StII_1-30_ forms pores with a toroidal shape structure, where the head groups of the lipids contribute to the pore architecture (18). Strikingly, peptide molecules were located on one edge of the pore (Figure 2B), which is enough to maintain the open configuration of toroidal pores, as previously reported for AMPs and for Bax (57). In the binary mixture, pore-opening was a one-step process that started after 300 ns with a rapid increase in pore size. In contrast, in POPE-containing membranes this was a multi-step process as reflected by the bimodal shape of the time course of the increase in the pore size (Figure 2E). Initially, a slow rise in the pore diameter from 0.5 to ~1.0 nm occurred after 250 ns, and later a rapid increase to 2.0 nm diameter took place until reaching stability (Figure 2E). This result suggests that POPE could catalyze the process of pore opening by StII_1-30_ in membranes. The final StII_1-30_ pore size in both systems was similar to the one previously experimentally estimated with the use of osmoprotectants of different sizes (9).

Figure 2F shows the variation of peptide inclination during the simulation time for both membrane systems. It was calculated as the mean value of the four peptide molecules forming pre-associated StII_1-30_, in each of the three individual replicas. In both cases, StII_1-30_ initially appeared parallel to the membrane surface but progressively tilted, adopting an oblique conformation with respect to the bilayer normal while embedding into the membrane. The maximal tilt angle in each system was reached around 250 ns, the same time in which stable pore opening started (Figure 2E). Figure 2G shows peptide orientation in the pore, calculated in the last 100 ns of simulation (when the open pore was stable (i.e. not change in the number of each lipid type surrounding the peptides and pore size). It was estimated as the mean value of the four peptide molecules forming associated StII_1-30_, in each individual replica. StII_1-30_ oriented such that the hydrophobic N-terminus of the peptide is buried into the pore-section, without spanning the membrane, while the C-terminus localizes at the water-membrane interface (Figure 2G). These results are also in line with the toroidal pore model in which peptide molecules do not need to completely penetrate the membrane but rather destabilize its structure by inducing membrane curvature.

### III.3 PE has a dual role in enhancing membrane association and functioning as a structural component of the pore formed by StII_1-30_

The comparison of the simulations in the membranes with and without PE allowed us to get insight into the role of this lipid in the mechanism of pore-formation of StII_1-30_. We found that PE enhanced both StII_1-30_ association to the membrane and pore formation (Figure 2C-E and Table 2). Even though the final pore size reached upon equilibrium was similar in all the membrane systems (Ø~2 nm), the kinetics of pore opening was faster in POPC:SM:POPE membranes (Figure 2E and Table 2). These results indicate that PE plays a key role in the mechanism of action of StII_1-30_ at facilitating the initial binding to the membrane and consequently the kinetics of pore opening.

In order to get further insight into the mechanism of how PE facilitates pore-formation by StII_1-30_, we compared the distribution of the different membrane lipids in the pore (Figure 3). Figure 3A shows the distribution of individual lipids in POPC:SM and POPC:SM:POPE membranes containing StII_1-30_ pores. Since this peptide induces membrane curvature during pore-formation, lipids can accommodate into the pore region according to their geometric properties. While POPC and SM are lipids with the propensity to form bilayer structures, POPE has a high tendency to adopt negative curvature due to its cone shape (55, 58) (Figure 3B). The latter class of lipids facilitates toroidal pore formation, in which the membrane bends to adopt a positive curvature in the direction parallel to the pore axis, and a negative curvature is defined at the pore boundaries (Figure 3C) (53, 55).

**Figure 3.**
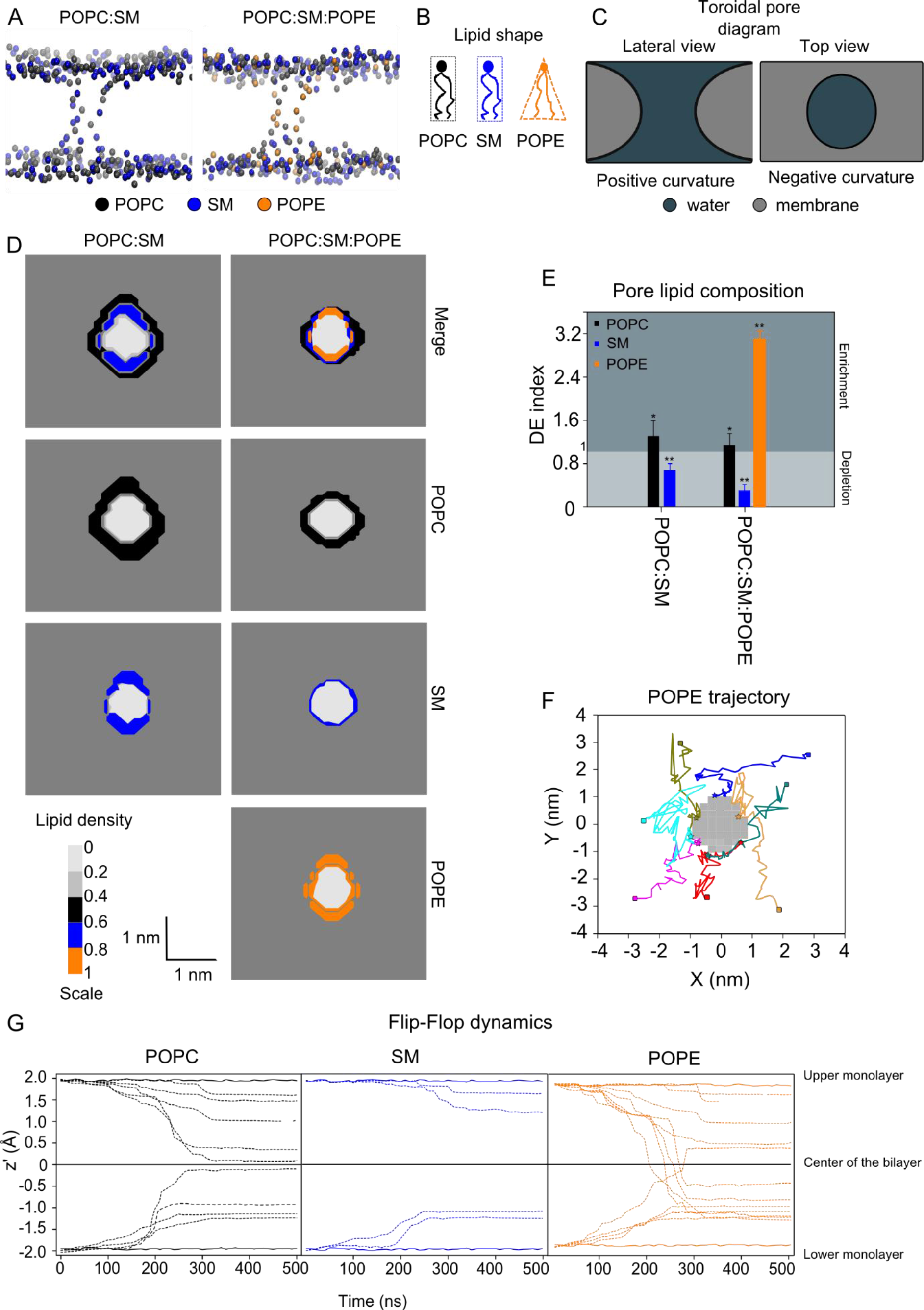
StII_1-30_ pores are enriched in POPE and POPC rather than SM. **(A)** Representation of the lipid composition of the toroidal pores induced by pre-associated StII_1-30_ after 500 ns simulation. **(B)** Shape of the lipids determines the propensity of the membrane to adopt planar or curved geometry. **(C)** Contribution of membrane curvature to the toroidal pore structure. Toroidal pore structures have positive curvature in the direction of the membrane bending and negative curvature around the ring structure. In relatively small pores, negative curvature has a dominant contribution. **(D)** Profile density of the lipids around the pore. **(E)** Depletion-enrichment (DE) index of the lipids during pore formation. The index was obtained from the last 100 ns of simulation out of 500 ns long simulation. Bars represent the average obtained from three replicas and error bars represent their standard deviation. Values higher than 1 indicate enrichment and values lower than 1 indicate depletion. Differences with p values ≥ 0.05 are indicated with * and < 0.05 with **. **(F)** A top-down view of the pore in the membrane, in which trajectories of individual POPE molecules during the 500 ns of simulation are shown. Each molecule is represented with different colors, initial position (□), final position (☆). **(G)** Dynamics of the flip-flop movement of lipids in POPC:SM:POPE (40:50:10) membrane, represented as the mean z coordinate profile for the head group beads (PO_4_) plotted over the 500 ns of simulation. The z coordinate was calculated over windows of 10 ns. Individual head group (**---**), average of the head groups located at the interface of the upper and lower monolayer (**▬**).

To analyze the lipid composition of the pores formed by StII_1-30_ in the different membrane systems, we calculated their lipid density profiles (Figure 3D). In all the studied compositions, specific lipid clustering was observed in the membrane region surrounding the pore (Figure 3D). The depletion-enrichment (DE) index allowed us to characterize the distinctive nature of the lipid environment surrounding the pore. This parameter quantifies the distribution of the individual lipids (POPC, SM or POPE) in the pore relative to the bulk of lipids in the membrane (Figure 3E). DE values higher than 1 evidenced enrichment and those lower than 1 showed depletion. On one hand, this analysis revealed a slight enrichment of POPC in the pore formed by StII_1-30_ in POPC:SM membranes. On the other hand, pores formed in POPC:SM:POPE presented a significant enrichment of POPE. In fact, even though the POPC:SM:POPE membranes contained just 10% of POPE, this lipid was mostly confined to the pore in this system (Figure 3E). This lipid was initially distributed along the bilayer but freely diffused from the membrane to the pore during 500 ns of simulation, as evidenced by the trajectories of individual POPE molecules (Figure 3F).

One relevant feature of toroidal pores is that membrane bending and curvature induction facilitate flip-flop movement of the lipids through the pore ring (59, 60). To compare the contribution of each individual lipid to the flip-flop movement in the process of pore formation, we calculated the *z* coordinates of those lipids involved in the pore structure (within a 1.5 nm cut-off from the center of the pore, considering that the final size of the pore is 2 nm) for POPC:SM:POPE. Figure 3G shows the position of the lipids forming the pore as a function of the simulation time. A coordinate of z=0 indicates the middle of the membrane, while +2 nm and −2 nm are the two interfaces with water on either side of the membrane. POPE and POPC showed more flip-flop events than SM. Altogether our analysis suggests that in contrast to the full-length toxin, SM is not a critical component of the pore formed by StII_1-30_ and that in the ternary composition the pore structure is mainly formed by POPE and POPC molecules. Among these lipids, POPE seems to play a critical role at facilitating pore formation due to its ability to form non-lamellar structures, which could assist membrane curvature induction during the assembly of toroidal pore structures and the consequent flip-flop movement of lipids.

### III.4 StII_1-30_ activity in membrane systems is enhanced by PE

From our simulations, PE was predicted as a relevant component of the membrane that could facilitate StII_1-30_ activity and as a structural element of the pore. This lipid has been classically considered an inductor of negative curvature in the membrane due to the structural relation tail > head (Figure 3B) (58). In order to confirm the in silico results, we studied the pore-forming mechanism of StII_1-30_ with experimental biophysical tools. We assessed the StII_1-30_ binding to lipid monolayers and liposome vesicles and evaluated its permeabilizing activity in the later system. Membrane systems were composed of PC:SM (50:50) or PC:SM:POPE (50:40:10) to evaluate the contribution of PE to peptide membranotropic activity.

The increase in surface pressure (Δπ) by the association of peptides with previously formed lipid monolayers at the air–water interface can be employed to characterize their ability to interact with organized lipids (17, 48, 61). This method has the advantage that it does not require fluorescent amino acids or probes to detect peptide-lipid interactions, thus complementing fluorescence based lipid binding assays (62). Figure 4A shows the kinetics of surface pressure (π) increase upon StII_1-30_ binding to a PC:SM:PE lipid monolayer. The injection of the peptide into the sub-phase of the monolayer triggered an increase in π (Δπ), which stabilized after ~100 seconds (Figure 4A). As expected, Δπ decreased with the increase of initial surface pressure (π_0_) due to tighter packing of the lipids (Figure 4B). A suitable parameter for the characterization of lipid-peptide interaction is the critical pressure (π_c_), obtained by extrapolating the Δπ at equilibrium as a function of π_0_ to zero (63, 64) (Table 3). This parameter corresponds to the minimum pressure that must be applied to avoid incorporation of the peptide into a monolayer and is directly correlated with its affinity for the monolayer. StII_1-30_ attained the highest π_c_ values in the PE-containing membrane, indicating that it probably penetrates more deeply into this lipid monolayer when compared with POPC:SM monolayers (Table 3).

**Figure 4.**
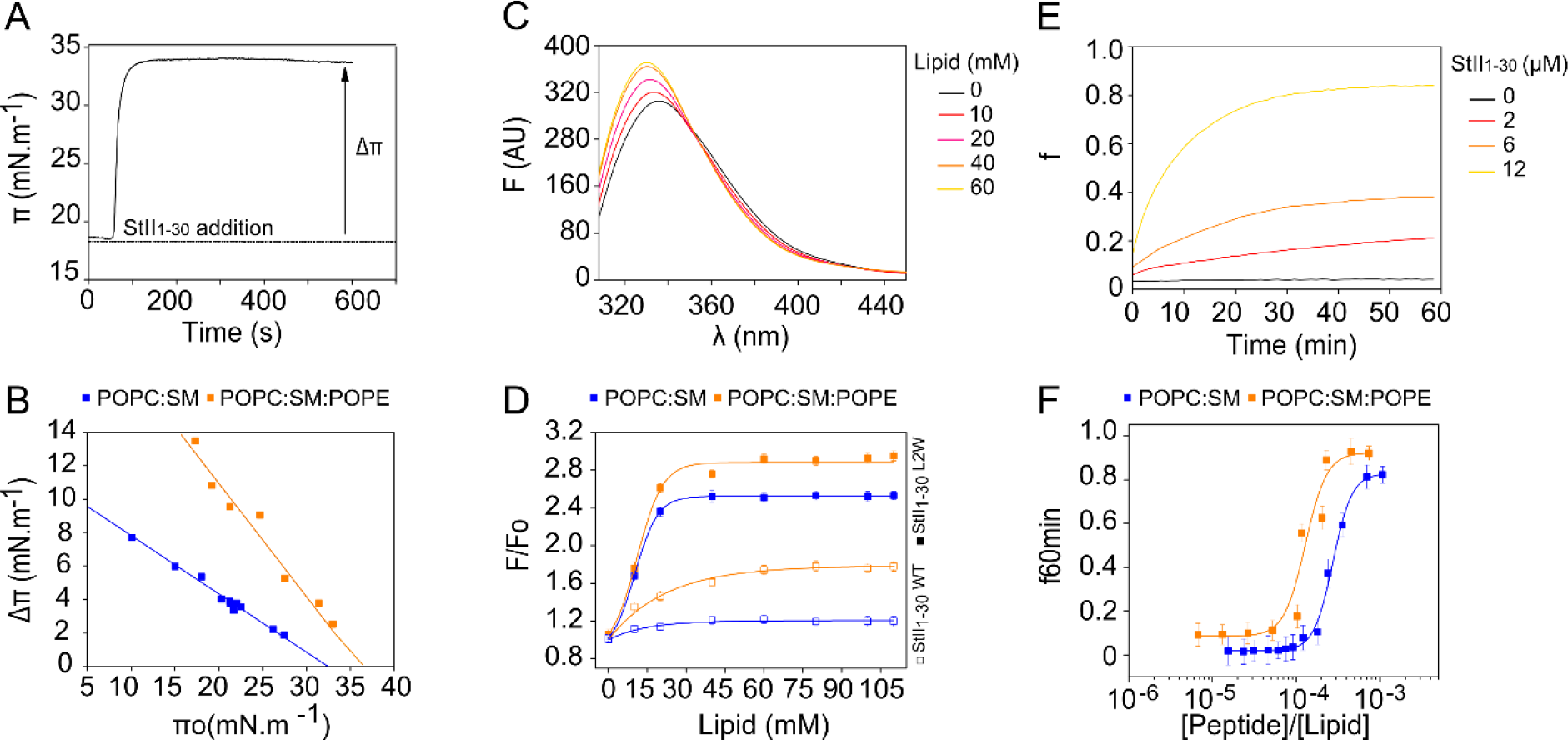
POPE favours binding to lipids and membrane permeabilization by StII_1-30_. **(A)** Kinetics of StII_1-30_ binding to a POPC:SM:PE (50:50) lipid monolayer. **(B)** Critical pressure induced by StII_1-30_ on monolayers of different lipid composition. Lines represent the best linear fit of the Δπ as a function of the initial monolayer pressure (π_o_), r^2^> 0.99. Peptide concentration: 1μM. **(C)** Fluorescence spectra of StII_1-30L2W_ in the presence of increasing amount of PC:SM:PE SUV. **(D)** Increase influorescence intensity as a function of lipid concentration. F_0_ and F stand for the fluorescence intensities (at λ_max_) from the peptide’s spectra in the absence and presence of SUV, respectively. λ_exc_: StII_1-30__(Phe14) 251 nm, λ_exc_: StII_1-30L2W__(Trp2) 280 nm, peptide concentration: 20 μM. **(E)** Kinetics of liposomes permeabilization induced by different concentrations of StII_1-30_ in PC:SM LUV **(F)** Release of CF from LUV promoted by StII_1-30_ after 60 min of assay. Lipid concentration: 10 μM, lipid compositions: PC:SM (50:50) or PC:SM:PE (50:40:10).

**Table 3:** Parameters that characterize StII_1-30_ activity in lipid model membranes.

| Parameters | Membrane |  |  |  |
| --- | --- | --- | --- | --- |
|  | POPC:SM |  | POPC:SM:PE |  |
|  | StII<sub>1-30</sub> WT | StII<sub>1-30</sub> L2W | StII<sub>1-30</sub> WT | StII<sub>1-30</sub> L2W |
| $\pi_c^a$ | 32.4 ± 0.6 | ND | 35.7 ± 0.4* | ND |
| F/F<sub>0</sub> max<sup>b</sup> | 1.20 ± 0.02 | 2.52 ± 0.02 | 1.78 ± 0.01* | 2.88 ± 0.02 * |
| Lip<sub>50</sub><sup>c</sup> | 14.5 ± 0.3 | 4.5 ± 0.1 | 22.2 ± 0.2* | 5.0 ± 0.2* |
| C<sub>50</sub> × 10<sup>4</sup><sup>d</sup> (mol:mol) | 3.2 ± 0.1* | ND | 0.9 ± 0.1 | ND |

<sup>a</sup>Critical pressure ( $\pi_c$ ) which corresponds to the pressure that must be applied to avoid incorporation of the peptide to the monolayer and is directly correlated with its affinity for the lipids. <sup>b</sup>Highest fluorescence intensity ratio (F/F<sub>0</sub>) obtained upon saturation conditions. <sup>c</sup>Amount of lipid necessary to bind half of the peptide molecules (Lip<sub>50</sub>) estimated from the Boltzmann function. F<sub>0</sub> and F are the fluorescence in the presence and absence of vesicles, respectively. <sup>d</sup>C<sub>50</sub> is the peptide concentration necessary to achieve 50% of vesicles permeabilization. Values correspond to the mean and standard deviation from at least three independent experiments. \*indicates that the values are statically different with p values calculated from a two-tailed distribution reported with a confidence level of 0.05.

Binding of StII_1-30_ to liposomes was measured by following the changes in Phe (native StII_1-30_) or Trp (StII_1-30L2W_) fluorescence emission spectra elicited by mixing with lipid vesicles (65). Figure 4C shows typical spectra obtained upon StII_1-30L2W_ titration with PC:SM:PE SUV. The addition of increasing amounts of liposomes to the peptide solution triggered a progressive increase in fluorescence emission intensity as well as a blue shift of the position of maximum emission (λ_max_), indicating a change in the Trp local environment (Figure 4C). The magnitude of the change determined at high lipid concentration is measured by the parameters [F/F_0_]_max_. Vesicles containing PE showed higher values of [F/F_0_]_max_ suggesting that they were better membrane targets for StII_1-30_ (Figure 4D and Table 3). To characterize the affinity of StII_1-30_ to lipid vesicles, the L_50_ was determined. This parameter represents the lipid-protein molar ratio at which half of the peptide molecules are bound to the liposomes (Table 3). The lowest L_50_ also showed that StII_1-30_ had more affinity for PE-containing membranes.

Permeabilization of lipid vesicles has provided direct evidence for the pore-forming activity of StII_1-30_ (12, 13, 18). Here we examined whether PE affected StII_1-30_ activity. Figure 4E shows the time course of StII_1-30_–elicited activity in PC:SM liposomes. Addition of StII_1-30_ to a solution containing CF-loaded LUVs promoted the release of the dye at different rates and to different fractions in a concentration-dependent manner (Figure 4F). This peptide had higher activity in PC:SM:POPE than in PC:SM vesicles, which was reflected in the C_50_ parameter that represents the peptide concentration required to achieve permeabilization of 50% of the vesicles (Table 3). Combined, our experimental results qualitatively support the MD simulations in that PE favors StII_1-30_ pore formation, probably by acting as an important element of the pore due to its intrinsic capability to form non-lamellar structures.

### III.5 StII_1-30_ activity has typical functional features of toroidal pores

Toroidal pore structures have been difficult to directly image due to their intrinsic structural flexibility. In spite of these difficulties, curvature effects and lipid flip-flop movement have been considered indirect evidence of this pore mechanism (4). The rationale behind this assumption relies on the toroidal shape of these structures. This model postulates the bending of the inner and outer monolayers forming half a torus. Therefore, in this pore configuration, the membrane adopts both a positive curvature in the direction of the membrane bending and negative curvature in the direction of the ring structure (Figure 3C) (66, 67). Moreover, each pore behaves as a point of fusion between opposite monolayers, favoring the otherwise restricted transbilayer or flip-flop movement of lipids (58, 68).

We evaluated the ability of StII_1-30_ to promote an increase in the L/H_II_ phase transition temperature of DiPoPE, an approach that has been previously used to study the mechanism of toroidal pore formation of some AMPs (69, 70). Representative scans of DiPoPE in the presence or absence of StII_1-30_ were obtained by DSC (Figure 5A). Pure DiPoPE exhibited a thermotropic transition around 43° C that was shifted to a higher temperature when StII_1-30_ (52° C) was added to the liposomes (Figure 5B). An increase in Tm value for the L/H_II_ phase transition of the DiPoPE can be interpreted as evidence of the capability of StII_1-30_ to induce positive curvature in the membrane. By stabilizing positive membrane curvature, StII_1-30_ promotes membrane stress in the opposite direction regarding curvature. Therefore, StII_1-30_ pores would essentially need to flip inside out to form the H_II_ phase, a process that requires more energy, as evidenced by the higher Tm.

**Figure 5.**
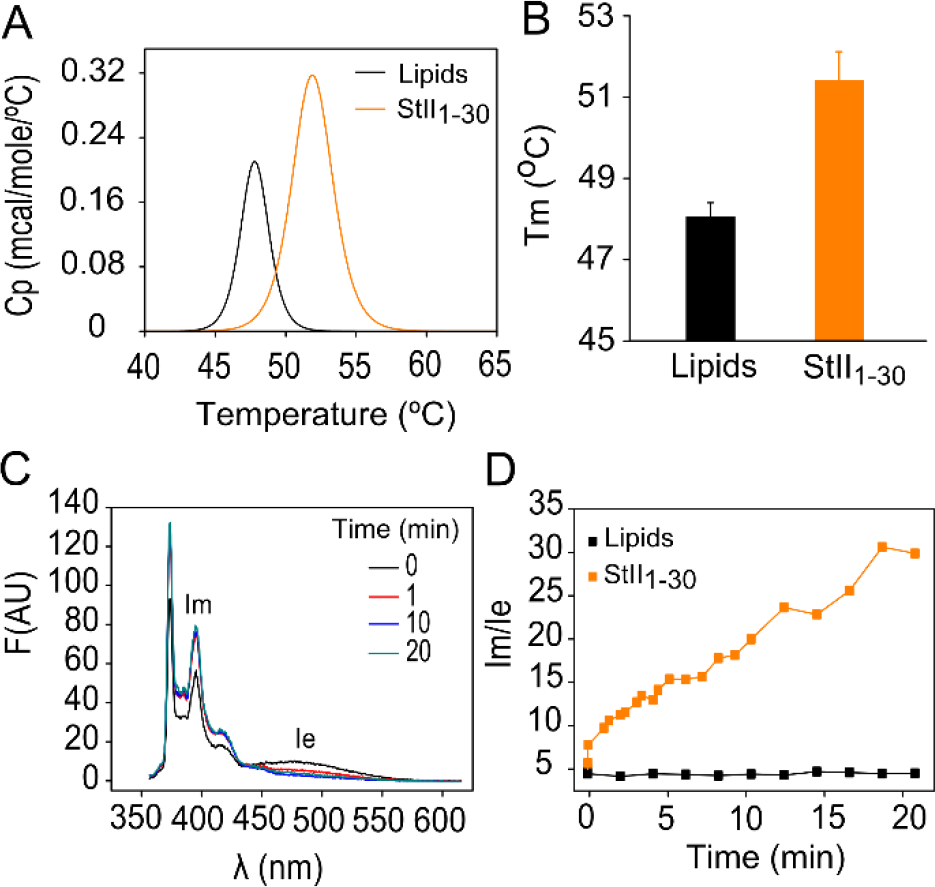
StII_1-30_ stabilizes positive curvature and induces flip-flop movement of the lipids in the bilayer. **(A)** DSC heating thermogram of MLV of DiPoPE in the presence or the absence of StII_1-30_. **(B)** Effect of StII_1-30_ on the bilayer (L) to hexagonal phase (H_II_) transition temperature of DiPoPE. Lipid concentration: 10 mM. Peptide:lipid ratio: 1:100. **(C)** PC-Pyrene spectra after different time of addition of StII_1-30_. **(D)** Kinetics of transbilayer redistribution of pyPC in the presence of StII_1-30_. The time course of the transbilayer redistribution of pyPC was followed by measuring the fluorescence intensity of pyrene monomers (I_M_) and excimers (I_E_) at λ 395 and 465 nm, respectively, and calculating the ratio I_E_/I_M_. Peptide concentration: 10 μM, lipid composition PC:SM:PE (50:40:10), lipid concentration: 20 μM.

Finally, we investigated if the formation of StII_1-30_ pores promotes flip-flop movement of the lipids, as previously demonstrated for sticholysins (51) and for their shorter versions (71). We measured the transbilayer movement of a pyrene-labeled analog of PC (pyPC), in PC:SM:POPE (50:45:10) lipid vesicles. The shape of the emission spectra of the pyrene depends on the contribution of two populations made up by monomers and dimeric excimers. The extent of excimer formation depends on the collision frequency of pyrene-labeled analogs. As a consequence of flip-flop movement of lipids, initially asymmetric pyPC molecules change their fluorescent properties due to redistribution between the membrane leaflets (51). Toroidal pore formation should facilitate flip-flop movement of the lipids, inducing the redistribution of pyPC molecules in both leaflets (72). StII_1-30_ addition provoked a decrease of the excimer fluorescence intensities, a clear indication of the dilution of pyPC molecules between both leaflets (Figure 5C). StII_1-30_ induced lipid flip-flop in PC:SM:PE, as demonstrated by the increase of the I_M_/I_E_ ratio over time (Figure 5D). I_M_/I_E_ did not change in the absence of the peptide, indicating the stability of the liposome system. These experiments allowed us to conclude that similar to what has been described for sticholysins (51), StII_1-30_ is able to induce flip-flop movement of the lipids. Given the ability of this peptide to form pores in model (12) and cellular membranes (9) this evidence supports a model in which it forms toroidal structures involving membrane bending and lipid diffusion through the pore.

## IV. Discussion

Sticholysins are typical eukaryotic α-PFTs that form pores upon insertion of their N-terminal lytic fragment into the membrane (73). They act through the formation of protein-lipid pores and have been extensively used as a tool to investigate the mechanism of toroidal pore formation in membranes (4, 5, 21, 22). Synthetic peptides corresponding to the N-terminus of sticholysins are good models for the study of the structure and function of these toxins (9, 10, 12, 13). StII_1-30_ is the most active peptide derived from the N-terminus of actinoporins (13), and therefore may be a promising candidate for biomedical applications. Here we combined MD simulations and biophysical tools to get further insight into the pore-forming mechanism of StII_1-30_. Specifically, we investigated the role of membrane curvature and lipid flip-flop on the mechanism of pore formation by this peptide. The combination of computational and experimental biophysical tools allowed us to gain information about the structure of the pore formed by StII_1-30_ in lipid membranes and the role of curved lipids in the assembly of toroidal pores.

We have previously proposed that the higher permeabilizing activity of StII_1-30_ could be ascribed to its higher propensity to fold and pre-associate as coiled-coil structures in solution, without the assistance of the membrane (29). The results of this work support the hypothesis that pre-association in solution is an essential prerequisite for the pore-forming activity of StII_1-30_. From our simulations, this peptide was unable to form pores independently of the lipid composition when bound to the lipid membrane in its monomeric form. Peptide association in solution seems to be a common requirement to form toroidal pores by peptides (74). As for StII_1-30,_ we propose that pre-association in solution might assist irreversible binding to lipids by increasing the local membrane-bound peptide concentration. The asymmetric attack of associated StII_1-30_ to one leaflet of the bilayer and its high local concentration at the membrane would facilitate bilayer destabilization and pore opening due to the increase of membrane tension, as proposed for other membranotropic peptides such as those derived from Bax (8, 75, 76).

In all the membrane systems containing pre-associated StII_1-30_, we observed the same sequence of events including: *i*) binding to the membrane interface, *ii*) enhancement of membrane thickness around the pore region, *iii*) curvature induction with the consequent membrane bending, and *iv*) pore formation. StII_1-30_ formed pores of approximately 2 nm diameter size, which mirrors our previous experimental data in human red blood cells (9). Pores formed by StII_1-30_ displayed typical features of toroidal pores, as also suggested for the full-length protein (15, 77). Peptide molecules appear accommodated in the membrane in an oblique configuration with the N-terminus nearer the core of the ring and the C-terminus laying in the membrane interface, in agreement with previous data obtained in liposomes (13, 28). A configuration in which StII_1-30_ is located at the pore surface without spanning the membrane bilayer and only in one edge of the ring, also supports the toroidal pore model. In this model, one section of the pore can remain protein-free, as observed for other proteins such as the cholesterol-dependent cytolysins and Bax or some AMPs (21, 22).

In the context of toroidal lipid pores, curvature induction represents a critical step for pore opening (4). In this case, the role of the peptides or lytic fragments is to help to reduce the stress caused by membrane distortion and stabilize curvature formation (67). Experimentally, StII_1-30_ induced positive curvature and flip-flop movement of the lipids, similarly to its derived form (15, 51, 77). Mechanistically, this could be explained based on the hydrophobic effect. Initial membrane disruption due to peptide binding can promote temporary exposure of the hydrophobic acyl chains of the lipids to the aqueous environment. To avoid the high energy cost of this process, lipids bend and form a highly curved, non-bilayer structure at the pore edge connecting the opposite monolayers of the membrane and forming a continuous surface. Afterward, lipids can easily exchange monolayers by simple diffusion over the pore edges. As a result, in the framework of this model, not only peptide molecules but also membrane mechanical properties play an important role in pore assembly (44, 60, 67).

Both computational and experimental approaches showed that membrane binding and pore formation by StII_1-30_ was enhanced by PE. The presence of this lipid in the membrane bilayer seems to facilitate pore formation by acting at two different stages of the process. On the one hand, PE could be considered a modulator of the membrane properties that are required for efficient pore formation due to its capability to form non-lamellar lipid phases (66, 78, 79). This capability seems to favor early stages of the membrane destabilization process by enhancing peptide insertion into the bilayer and membrane remodeling events, which are fundamental to accomplish stable pore opening. Given that the generation of a pore is energetically expensive, the presence of such non-lamellar lipids contributes to pore stability (80). On the other hand, PE appears as a co-factor of the pore induced by StII_1-30,_ since membrane regions comprising the pore assembly are especially enriched in this lipid. It is well-known that the pore size strongly influences the balance of the contribution between positive and negative curvature in toroidal pores (54). Positive curvature is induced in the direction of the membrane bending and is modulated by the membrane thickness, while negative curvature is stabilized in the direction of the ring, parallel to the membrane (54). Knowing that curvature is inversely proportional to the radius, the smaller the pore size is, the higher the negative curvature along the pore edges. As a consequence, in the case of very small pores, like those formed by actinoporins (15, 81) or StII_1-30_ (9), negative curvature-inducing lipids (i.e PE) could easily accommodate in the ring sections of the pore and play an additional role at stabilizing its edges (Figure 4C).

## Conclusions

We propose a mechanistic framework that explains in detail the lytic activity of StII_1-30_ and provide evidence for a toroidal model for the structure of the pore it forms. Membrane curvature induction and induction of flip-flop movement of the lipids were identified as two important membrane remodeling steps related to toroidal pore formation by StII_1-30._ We highlight the key role of PE and in general curved-shape lipids as co-factors of toroidal pores that can facilitate the adoption of the open state by easily accommodating in the pore ring. Understanding these aspects of the mode of action of StII_1-30_ could underpin strategies for the rational use of this and other peptides or proteins acting via toroidal pore formation as biotechnological tools in different lipid formulations.

## Author contributions

HMG performed and analyzed the MD simulations, the fluorescence assays and the flip-flop experiments. UR performed and analyzed the calorimetry and the monolayer experiments. PV and DPT contributed to MD simulation design. RFE and RME supervised the calorimetry experiments and provided resources. CA and ML contributed to the design of the biophysics experiments and contributed the conception of the project. CA, DPT and UR supervised the work and provided resources. HMG, DPT and UR wrote the manuscript with input from other authors. DPT and UR designed and led the project.

## Acknowledgements

UR was grantee from IFS (4616). DPT holds the Alberta InnovatesTechnology Futures Strategic Chair in (Bio)Molecular Simulation. HMG is an Alberta Innovates Health Solutions Scholar and Vanier Canada Research scholarship recipient. Computational resources were provided by Compute Canada, funded by the Canada Foundation for Innovation and partners. Additional support came from the Canada Research Chairs Program (DPT) and the Canadian Institutes of Health Research (DPT). Authors thank Dr. Katia Cosentino, Dr. Rodrigo Cuevas and Dr. Lohans Pedrera for carefully reading the manuscript.

